# Profiling IOP-responsive genes in anterior and posterior ocular tissues in the rat CEI glaucoma model

**DOI:** 10.1101/2024.02.11.579818

**Authors:** Diana C. Lozano, Yong-Feng Yang, William O. Cepurna, Barbara F. Smoody, Eliesa Ing, John C. Morrison, Kate E. Keller

**Affiliations:** Casey Eye Institute, Oregon Health & Science University, 3181 SW Sam Jackson Park Road, Portland, OR 97239

**Keywords:** gene expression, trabecular meshwork, optic nerve head, intraocular pressure, glaucoma

## Abstract

**Purpose:** The rat Controlled Elevation of Intraocular pressure (CEI) model allows study of *in vivo* responses to defined intraocular pressures (IOP). In this study, we use Nanostring technology to investigate *in vivo* IOP-related gene responses in the trabecular meshwork (TM) and optic nerve head (ONH) simultaneously from the same animals.

**Methods:** Male and female rats (N=35) were subject to CEI for 8-hours at pressures simulating mean, daytime normotensive rat IOP (CEI-20), or 2.5x IOP (CEI-50). Naïve animals, receiving no anesthesia or surgical interventions, served as controls. Immediately after CEI, TM and ONH tissues were dissected, RNA isolated, and samples were analyzed with a Nanostring panel containing 770 genes. Post-processing, raw count data were uploaded to Rosalind® for differential gene expression analyses.

**Results:** For the TM, 45 IOP-related genes were significant in the “CEI-50 vs. CEI-20” and “CEI-50 vs. naïve” comparisons, with 15 genes common to both comparisons. Bioinformatics analysis identified Notch and TGFβ pathways to be the most up- and down-regulated KEGG pathways, respectively. For ONH, 22 significantly regulated genes were identified in the “CEI-50 vs. naïve” comparison. Pathway analysis identified ‘defense response’ and ‘immune response’ as two significantly upregulated biological process pathways.

**Conclusions:** This study demonstrates the ability to assay IOP-responsive genes in both TM and ONH tissues simultaneously. In the TM, downregulation of TGFβ pathway genes suggest that TM responses may prevent TGFβ-induced extracellular matrix synthesis. For ONH, the initial response to elevated IOP may be protective, with astrocytes playing a key role in these gene responses.

## Introduction

Intraocular pressure (IOP) is regulated by the trabecular meshwork (TM), a tissue that is located in the angle of the anterior chamber. Normotensive pressures are required to maintain the shape of the eye globe and allow consistent focus of light on the retina. Elevated IOP occurs due to blockage of the aqueous outflow channels in the TM and is a major risk factor for glaucoma. ^1^ TM cells detect IOP elevation and in response, they alter gene transcription to homeostatically adjust IOP back to normal ranges. ^1–3^ With IOP elevation, retinal ganglion cells die, likely due to initial injury of their axons at the level of the optic nerve head (ONH). ^4, 5^ Identifying which genes respond to elevated IOP will be important to develop new IOP-lowering therapies that protect the TM and ONH.

Different approaches have been used to study the effects of elevated IOP on TM gene expression. Static or cyclic mechanical stretch of cultured TM cells produces an approximately 10% stretch, mimicking what TM cells experience when IOP is elevated *in vivo*. ^6–8^ This model has been used in many microarray and qPCR studies to evaluate which genes are likely to respond to elevated IOP. However, the main criticism of this technique is that the TM cells are taken out of their native environment and are grown on a plastic or silicone membrane. *Ex vivo* perfusion of anterior segments is also commonly used. ^2, 9–13^ In this organ culture method, the whole globe is bisected, the iris, ciliary body, and lens are removed, and the resulting anterior segment, containing cornea, TM and sclera, is clamped into a perfusion apparatus. ^9^ Serum-free media is perfused either at 1x pressure (8 mmHg) or 2x pressure (16 mmHg) to mimic the elevated IOP experienced by glaucoma patients. Constant flow systems are also used, where IOP is continuously monitored. However, because the globe is dissected, anterior segments are not subject to normal IOP fluctuations and aqueous dynamics found *in vivo*. Animal models are necessary to understand better the role of cellular responses to IOP in the living eye.

Numerous animal models of ocular hypertension are used in glaucoma research. These include the DBA/2J mouse and transgenic mouse models such as mice over-expressing the human Y437H myocilin mutation. ^14, 15^ IOP can also be elevated by intracameral injection of viral vectors over-expressing bioactive molecules such as transforming growth factor beta (TGFβ), ^16, 17^ gremlin, ^18^ secreted frizzled-related protein-1 (SFRP1), ^19^ Dickkopf-related protein-1 (DKK1), ^20^ or CD44. ^21^ Other methods to increase IOP are injection of polystyrene microbeads into the anterior chamber, injecting hypertonic saline into Schlemm’s canal via episcleral veins, or by laser photocoagulation of TM. ^22–25^ Most of these models were primarily developed to study mechanisms of pressure-induced optic nerve damage and generally compromise TM tissue structure and function. Thus, while they are useful for studying neuropathology, they cannot be used to study the effects of IOP on TM genes. For practical reasons, IOP is monitored sporadically in these models, generally only once or twice per week, which likely misses pressure spikes or IOP fluctuations. Thus, the actual IOP is not fully known, which can profoundly affect interpretation of results.

An additional animal model of IOP, Controlled Elevation of IOP (CEI), has been recently described. ^26, 27^ IOP elevation is achieved by cannulating the anterior chamber and raising a reservoir of fluid to deliver IOP at a known pressure for a desired duration. IOP is continuously monitored in these anesthetized CEI rats, along with several physiologic parameters. We recently used this model to demonstrate the sequence of ONH cellular responses following a single, 8 hour pressure stimulus, ^27^ and identified several genes and pathways in common between CEI and those previously reported in chronic models of ocular hypertension. ^27^ For example, we found an immediate upregulation in defense related genes and this was followed by upregulation in cell proliferation. Thus, the CEI model encapsulates cellular events that occur in chronic glaucoma, but with the added benefit of being able to identify sequential events that lead to axonal degeneration. With its ability to control the duration and extent of IOP elevation, this model can ultimately be used to test the effect of inhibition or enhancement of specific genes and pathways on axonal survival.

While the CEI model was specifically designed to evaluate ONH cellular events, we have now adapted it to help us also understand TM responses to discrete pressure insults. From the perspective of TM research, this is the only model that allows study of an undamaged TM exposed to elevated IOP. The cannula tip resides anterior to the iris and elevated IOP does not cause angle closure, but induces a stretch/distortion of TM cells similar to that found in hypertensive glaucoma patients. Furthermore, we have recently described a method to microdissect the TM from rat eyes and isolated sufficient RNA for gene expression analysis. ^28^ This produced relatively pure TM that was not substantially contaminated with ciliary body tissue, as defined by quantitative PCR using biomarkers of the ciliary body. With these advances we are now able to investigate IOP-related gene response in the TM in a live rodent model for the first time.

In this study, we performed CEI at 50 mmHg for 8 hours and used Nanostring technology to compare TM gene expression changes to two control groups. We also analyzed gene expression changes in the ONH, providing the first study to investigate, simultaneously, IOP-related changes in the two tissues that are primarily affected by glaucoma, within the same animal.

## Methods

### Animals

A total of 35 retired Brown Norway breeder rats (Charles River Laboratories, Wilmington, MA) were used for this study. They were 6-9 months old with an equal number of male and female animals, equally distributed between experimental groups. Rats were kept in a standard 12-hour light and 12-hour dark cycle with *ad libitum* access to food and water. Animal protocols were approved by the Institutional Animal Care and Use Committee at the Oregon Health & Science University, were performed in accordance with the National Research Council’s Guide for the Care and Use of Laboratory Animals, and adhered to the ARVO Statement for the Use of Animals in Ophthalmic and Vision Research.

### The Rat CEI Model

CEI was performed in anesthetized rats as previously described, with constant monitoring of body temperature, O_2_ saturation, heart rate, and blood pressure by MouseOx and CODA. ^26^ Briefly, one eye of each rat was cannulated with a 1” polyurethane tube (0.010” OD/ 0.005 ID”; Instech Laboratories Inc., Plymouth Meeting, PA). The tip of the cannula was positioned anterior to the iris and TM (Figure 1A), under microscopic visualization, to ensure that the results would be associated with IOP-related stretching of the TM, rather than angle closure and inflammatory responses from iris contact with the TM. The cannula was connected to larger tygon tubing (Component Supply Company, Fort Meade, FL) attached to a reservoir with sterile balanced salt solution plus (BSS+; Alcon Laboratories, Inc.) and a pressure transducer. The height of the reservoir determines the extent of pressure elevation. ^26^ Previous work has shown that rat daytime mean IOP is ∼20 mmHg. ^29–35^ Therefore, we chose 50 mmHg (CEI-50), which is 2.5x daytime IOP, for the pressure challenge group (n=12). This IOP is consistent with the extent of pressure challenge used in human *ex vivo* studies. ^2, 8, 13^ One control group received the normotensive daytime mean IOP of 20 mmHg (CEI-20) (n=11). An 8-hour CEI pressure challenge was chosen because >6-hours of pressure challenge is required to produce an IOP homeostatic response in *ex vivo* perfused human anterior segments. ^2, 3, 12, 36^ Importantly, we have found that animals can tolerate 8 hours of general anesthesia, with no measurable impact on their physiology. ^26^ A naïve control group of rats (n=12), with no anesthetic or surgical manipulations, was also included. During CEI, IOP was monitored every 30 minutes by TonoLab to detect leakage or cannula failure. Mean Tonolab IOP during CEI-20 and CEI-50 are presented in Figure 1B. To prevent eyes drying out, alternating applications of topical artificial tears or 0.5% proparacaine hydrochloride (Akorn, Inc, Lake Forest, IL) were applied every 15 minutes to both eyes. At the end of CEI, IOP was reduced to 20 mm Hg for 5 minutes, the cannula was removed, and the needle track self-sealed. Rats were then euthanized and their eyes immediately enucleated for TM and ONH microdissection.

**Figure 1.**
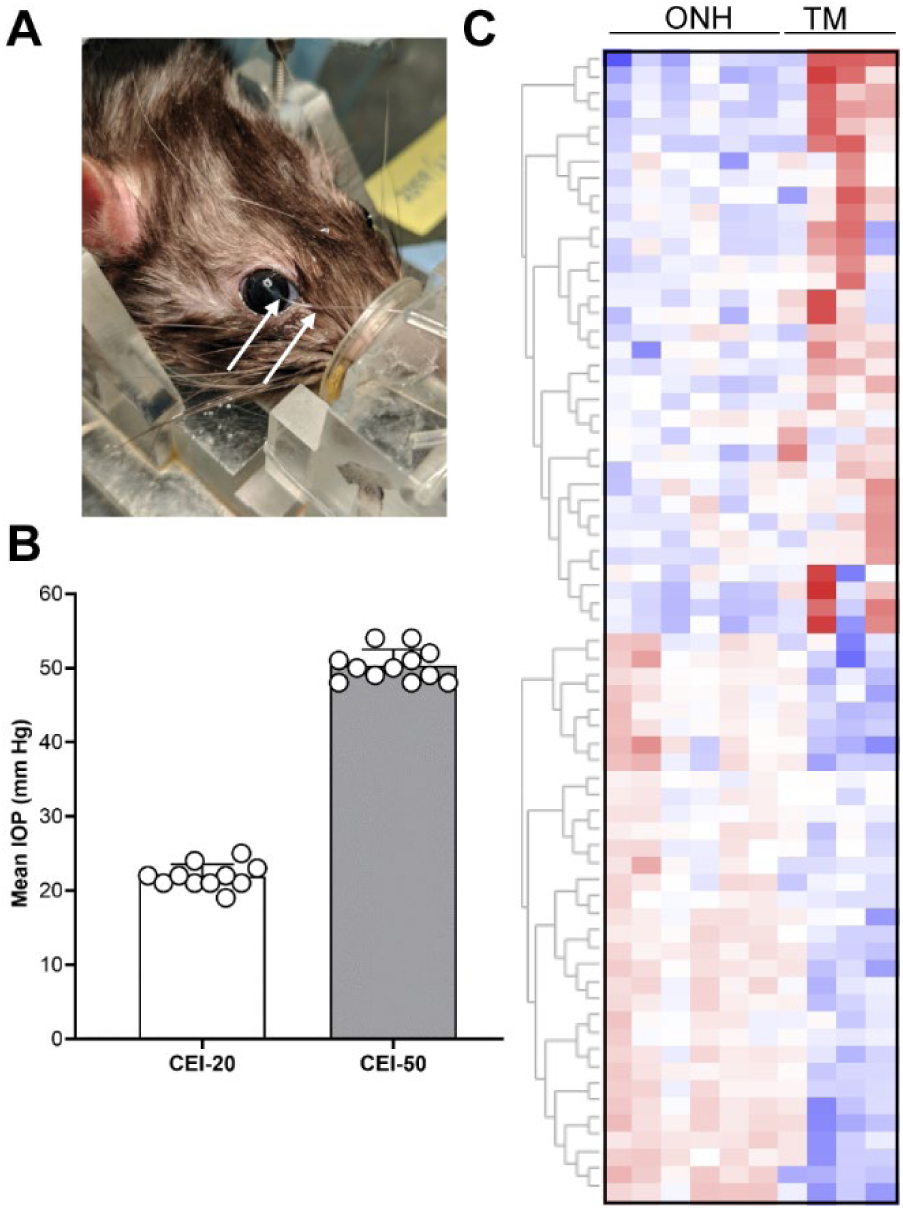
(A) Rat eye cannulation. Arrows point to the small tube inserted through the cornea, by which BSS+ fluid is delivered at a precise IOP. (B) Graph shows the mean IOP of each animal over the 8-hour duration of the experiment in the CEI-20 normotensive control group and the CEI-50 hypertensive experimental group. (C) Heatmap and dendrogram of ONH and TM genes from naïve samples showing differences in gene expression in the two tissues.

### TM and ONH microdissection and RNA isolation

TM and ONH dissection occurred as described in previous publications. ^27, 28, 37^ Following dissection, the tissues were placed in individual tubes with 200 ul TriZol and RNA was isolated as described. ^28^ The NanodropOne (ThermoScientific, Waltham, MA) was used to measure RNA concentration. RNA was isolated and mean RNA concentration was determined to be 61 ng/TM and 60 ng/ONH.

### RNA amplification and Reporter Probe Hybridization

Our samples had low RNA concentrations so the low RNA input reagent kit was used to amplify the target following the manufacturer’s protocol (Nanostring, Seattle, WA). Briefly, 4 µl of RNA was mixed with 1 µl RT master mix, placed in a thermocycler, and then converted to cDNA using the following protocol: 25 °C for 10 min, 42 °C for 60 min, 85°C for 5 min, and then placed on ice. A multiplexed target enrichment (MTE) was then performed for all samples as follows: Each cDNA sample was mixed with 1.5 µl 5X dT Amp master mix and 1 µl low input primers. Enrichment was performed with the following parameters: 95 °C for 10 min, 18 cycles of 95 °C for 15 secs and 60 °C for 4 min. To ensure successful RNA amplification, 1 µl MTE reaction was quantitated by Qubit 3.0 fluorometer analysis at the OHSU Gene profiling core facility. The results showed an average of 149.2 ± 3.74 ng/µl, ranging from 122-163.5 ng/µl (n=12). The remaining volume of MTE reactions were heated to 95 °C for 2 min, snap-cooled on ice, and mixed with 8 µl of hybridization master mix containing 5 µl hybridization buffer and 3 µl reporter probe codesets. After mixing, 2 µl capture probe sets in hybridization buffer were added to each sample. The reporter codesets contain half of the target-specific sequence and 6 fluorescent labelled RNA segments (barcodes), of 4 different colors. Unique combinations of these barcodes allow up to 972 target genes to be assayed at one time. The capture probe set contained the other half of the target-specific sequence and was labeled with biotin. The capture and reporter probes were hybridized to gene targets in each of our TM and ONH samples. Tubes were placed in a pre-heated 65 °C thermocycler for 24 hours. Hybridization reactions were then immediately transferred to the nCounter SPRINT profiler instrument.

### nCounter Sprint Profiler Analysis

Since predesigned rat-specific Nanostring panels are not available, the hybridized target-probe complexes were immobilized on *Mus musculus* PanCancer IO 360 cartridges (Nanostring). These contain 770 gene targets related to immune response (8 pathways), tumor microenvironment (4 pathways), and tumor biology (4 pathways). Internal reference genes are also included. Each cartridge can process 12 samples at one time and a total of 5 cartridges were used for these studies (see Supplemental Table S1). TM samples were randomized over 3 of these cartridges, while ONH samples were randomized over an additional 3 cartridges (i.e. 1 cartridge contained both TM and ONH samples). Cartridges were placed into the nCounter SPRINT Profiler instrument and hybridized samples were injected into the cartridge. The instrument follows a series of automated steps that include immobilization on a streptavidin-coated surface, flowing fluids over the surface to align the probes, and then data counting of the individual barcodes by a fluorescence microscope. Multiple fields of view are imaged and the number of fluorescent barcodes is counted. One barcode is equivalent to 1 RNA molecule.

### Rosalind and Bioinformatics Analysis

Raw data files (*.RCC) for each experiment were analyzed by Rosalind® informatics software. For TM, three comparisons were made: “CEI-50 vs. CEI-20”, “CEI-50 vs. naïve”, and “CEI-20 vs. naïve”. For ONH, only “CEI-50 vs. naïve” samples were compared since we found no significant gene responses in the CEI-20 vs. naïve comparison in our previous RNA-seq study. ^27^ Samples were excluded if they had low image quality (<0.8), high binding density (>1), or if housekeeping genes had zero counts. The final number of samples, derived from 27 rats, included in analyses were: CEI-50, n=6; CEI-20, n=8; and naïve, n=4 for TM samples; and CEI-50, n=5; and naïve, n=7 for ONH. Digital counts were normalized to ∼18 internal housekeeping genes on the panel. Background thresholds are calculated in the software as 97.5^th^ percentile of the negative controls and are removed from further analysis if more than half the samples do not meet this threshold. If this occurs, the log_2_ Fold Change is reported as 0 and the level of significance as 1. Differentially expressed genes were identified using a cutoff of ≥1.5 or ≤ -1.5 fold change and were considered significant if p-value was <0.05. Since RNA concentrations were low for each individual sample, technical replicates could not be performed. To investigate pathways, differentially expressed gene lists were uploaded to ShinyGO, version 0.77.

### Quantitative PCR

Taqman qPCR analysis was performed for a subset of the differentially expressed genes identified by Nanostring. Briefly, predesigned, rat-specific primers were purchased (Thermo Fisher Scientific), including primers for 18S and GAPDH, which were used as housekeeping genes for TM and ONH, respectively. Rat TM RNA was converted into cDNA using Superscript III reverse transcriptase (Thermo Fisher Scientific). cDNA was pre-amplified with pooled primers for the target genes. PCR was amplified using the following conditions: 95 °C for 10 min, 18 cycles of 95 °C for 15 secs and 60 °C for 4 min, and finished with 99 °C for 10 min. The pre-amplified cDNA was diluted 1:8, and 4.5 µl of this diluted sample was added in each PCR reaction. Quantitative PCR was performed on a quantstudio3 thermocycler (Applied Biosystem). Results were analyzed by Quantstudio 3 software, using a geometric mean to calculate ΔΔCt, and results were exported to Excel. Results were statistically analyzed using an unpaired t-test in Graphpad Prism 10.

## RESULTS

Nanostring technology was used to investigate *in vivo* IOP-related gene responses in the TM and ONH from the same animals. Initially, gene expression between TM and ONH samples from naïve animals we compared to validate that the technology was able to detect differences between the two naïve tissues. Not surprisingly, naïve TM samples clustered together and into a group distinct from naïve ONH samples (Figure 1C). These findings support the idea that different genes are regulated in naïve TM and ONH tissues. We will now address significantly regulated genes in the TM and ONH following CEI.

### TM gene analyses by Nanostring

Genes significantly altered in the “CEI-50 vs. CEI-20” comparison are shown in Table 1. Of 750 genes on the panel, 4 TM genes were significantly up-regulated and 26 were significantly down-regulated, as shown in the volcano plot in Figure 2A. The log2 normalized expression for the 4 up-regulated genes (*Cblc, Jag2, Pdzk1ip1*, *Prom1*) and 4 selected down-regulated genes (*Vegfa, Relb*, *Irf1, Inhba*) are also shown (Figure 2B). Pathway analysis of all 30 significantly regulated genes identified ‘defense response’, ‘immune response’, and ‘response to other organism’ as the top 3 significantly regulated ‘GO Biological Process’ pathways (Figure 2C).

**Figure 2.**
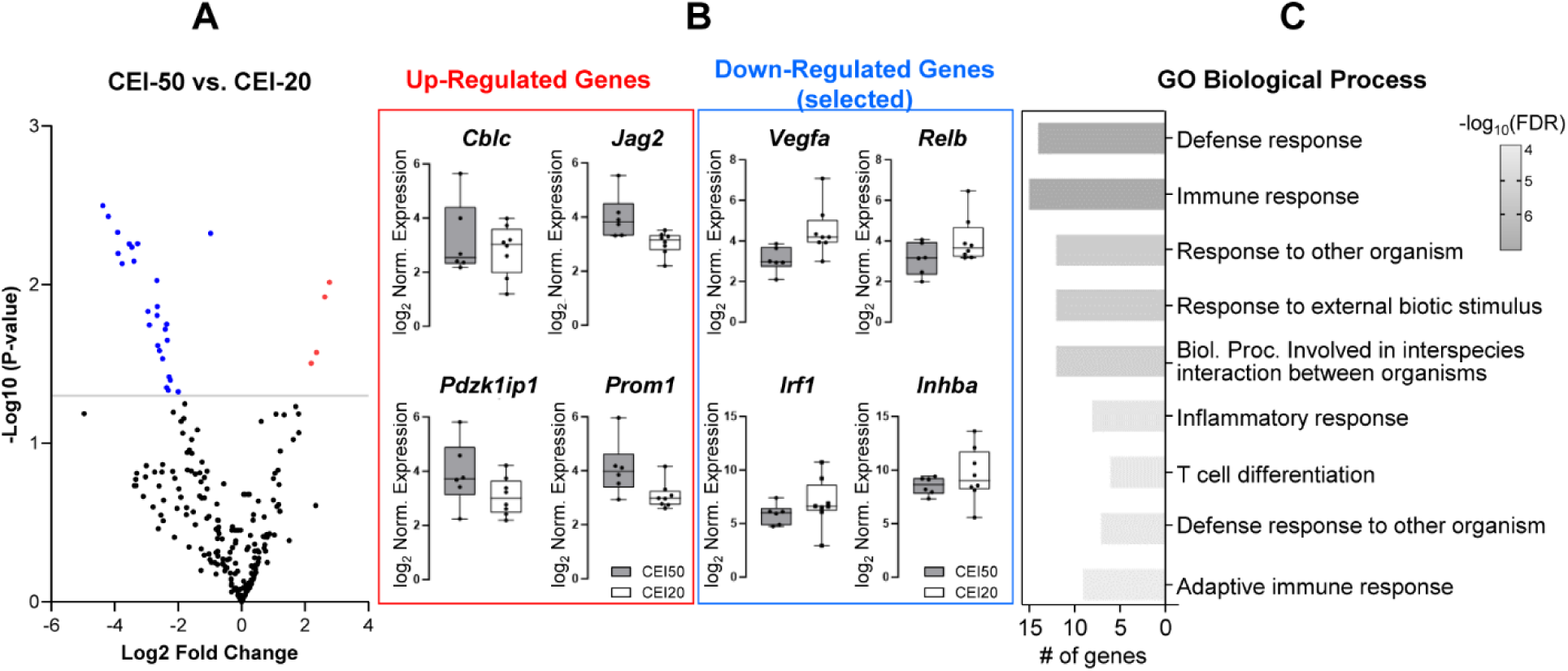
Nanostring analysis of CEI-50 versus CEI-20 groups. (A) Volcano plot showing all genes on the cartridge. Significantly up-regulated genes are shown in red and down-regulated genes are shown in blue. (B) Normalized expression levels of selected up- and down-regulated genes in CEI-50 (n=6; grey) and CEI-20 (n=8; white) are shown. (C) ShinyGO 0.77 analyses shows the top 10 most significantly affected ‘GO biological process’ pathways.

**Table 1:**
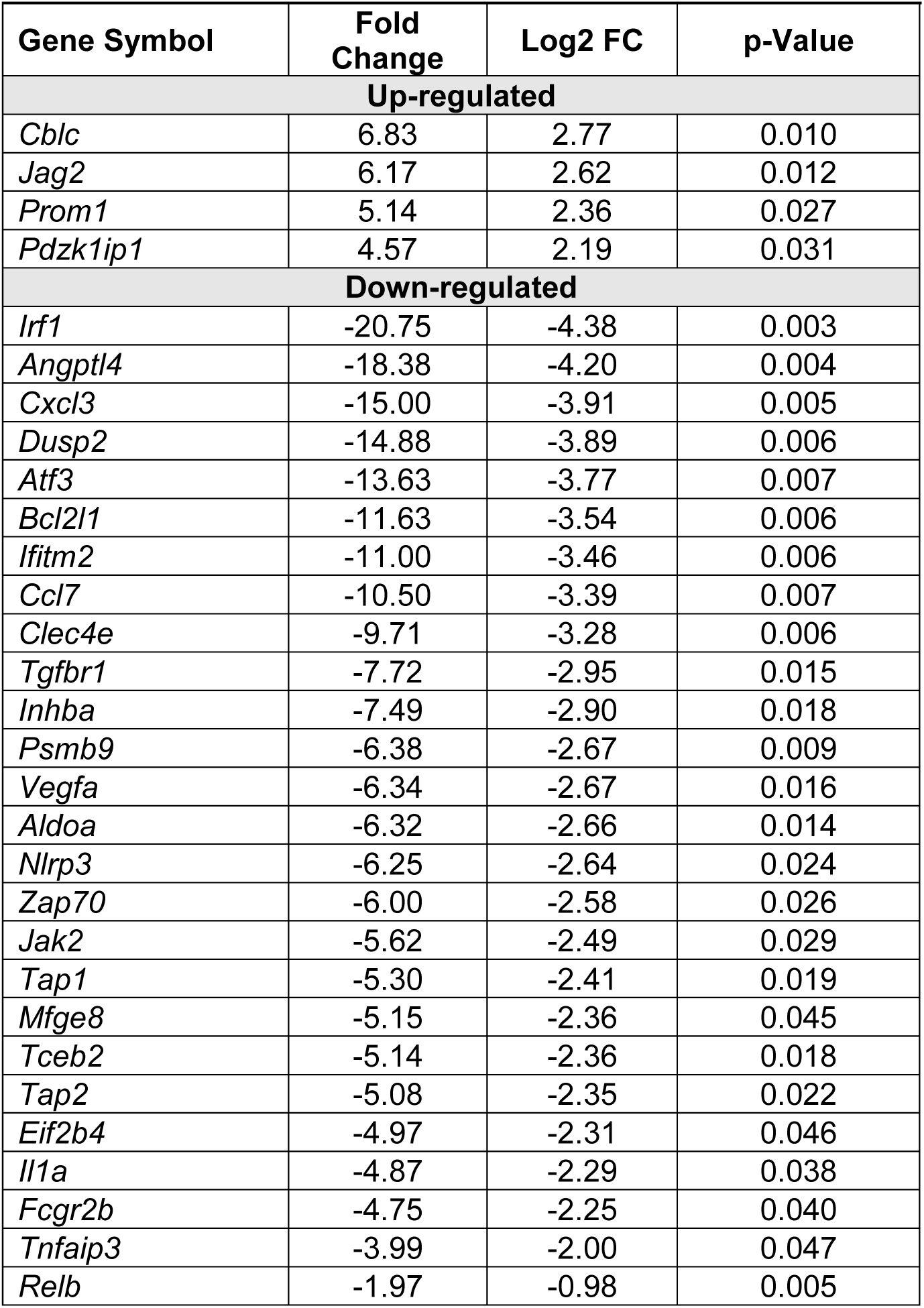
All TM genes significantly differentially expressed in “CEI-50 vs. CEI-20” (Fold change >1.5 or <-1.5; p-value <0.05).

In the “CEI-50 vs. naïve” comparison, there were 19 up-regulated and 63 significantly down-regulated TM genes, of which the top 20 are shown in Table 2 (the complete 63 gene list can be found in Supplemental Table S3). A volcano plot shows the distribution of all the genes in the “CEI-50 vs. naïve” comparison (Figure 3A). The log2 normalized expression for 4 randomly selected up-regulated (*Mmp13, Spry4, Stc1, Tnfaip6*) and down-regulated (*Aldoa, Mfge8, Rock1, Tgfbr1*) genes are also shown (Figure 3B). Pathway analysis of all 82 significantly regulated genes identified ‘regulation of cell population proliferation’, ‘cell population proliferation’, and ‘cell death’ as the top 3 of 10 significantly regulated ‘GO Biological Process’ pathways (Figure 3C). There were 111 significantly regulated genes in the two hypertensive comparisons (“CEI-50 vs. CEI-20” (n=30) and “CEI-50 vs. naïve” (n=81)), but 16 genes were identified in both comparisons (i.e. counted twice). Thus, there were 95 unique genes identified in our hypertensive comparisons. However, these genes may include those related to cannulation or anesthesia. To identify TM gene responses to elevated IOP alone, we next identified significantly regulated genes in the “CEI-20 vs. naïve” comparison.

**Figure 3.**
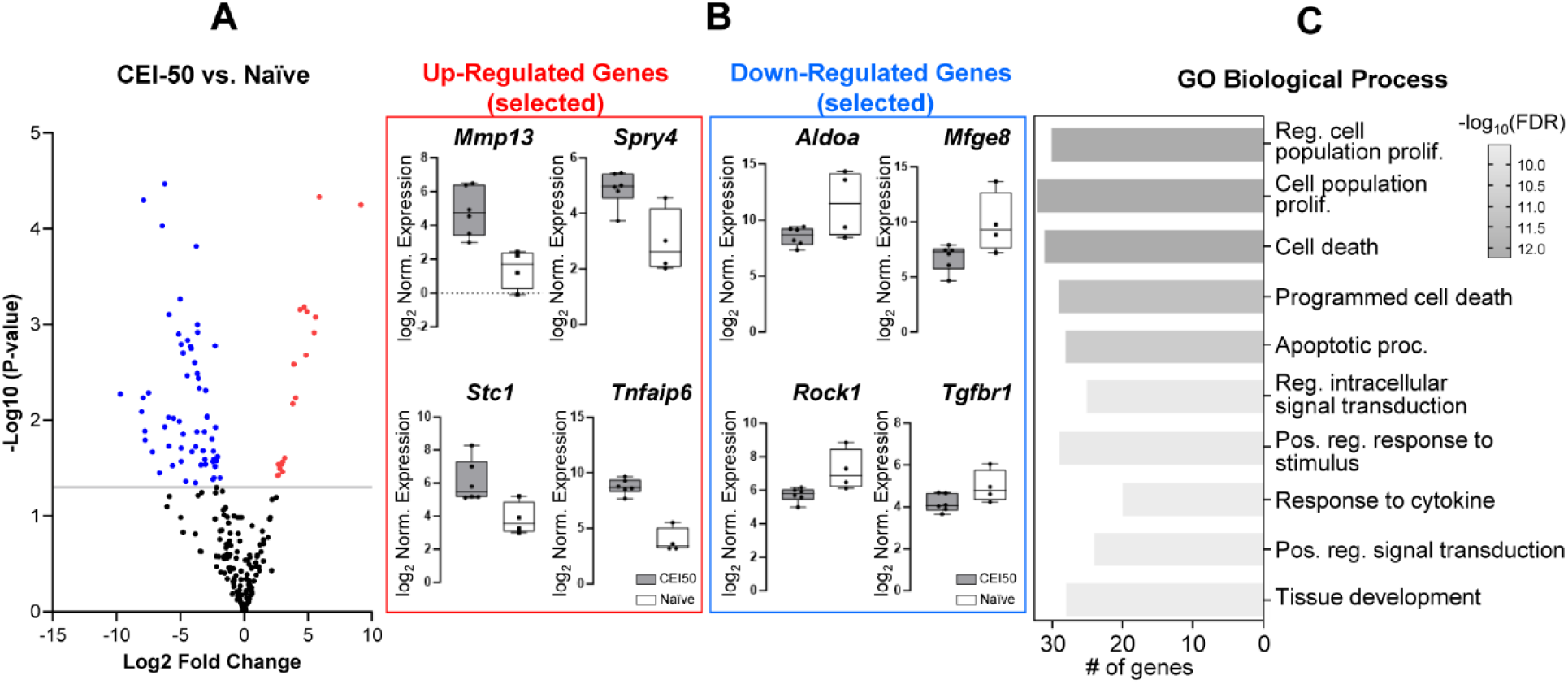
Nanostring analysis of CEI-50 versus naïve groups. (A) Volcano plot showing all genes on the cartridge. Significantly up-regulated genes are shown in red and down-regulated genes are shown in blue. (B) Normalized expression levels of selected up- and down-regulated genes in CEI-50 (n=6; grey) and naïve (n=4; white) are shown. (C) ShinyGO 0.77 analyses shows the top 10 most significantly affected ‘GO biological process’ pathways.

**Table 2:** All up-regulated and 20 down-regulated TM genes significantly differentially expressed in “CEI-50 vs. naïve” (Fold change >1.5 or <-1.5; p-value <0.05).

| Gene Symbol | Fold Change | Log2 FC | p-Value |
| --- | --- | --- | --- |
| **Up-regulated** |  |  |  |
| *Il6* | 565.09 | 9.14 | 5.63E-05 |
| *Tnfaip6* | 58.43 | 5.87 | 4.66E-05 |
| *Fosl1* | 47.83 | 5.58 | 0.001 |
| *Birc3* | 44.50 | 5.48 | 0.001 |
| *Nfil3* | 30.00 | 4.91 | 0.001 |
| *Mmp13* | 28.33 | 4.82 | 0.002 |
| *Myc* | 26.11 | 4.71 | 0.001 |
| *Cebpb* | 20.63 | 4.37 | 0.001 |
| *Ripk3* | 16.27 | 4.02 | 0.006 |
| *Stc1* | 14.76 | 3.88 | 0.003 |
| *Ccl2* | 14.05 | 3.81 | 0.007 |
| *Prom1* | 9.00 | 3.17 | 0.025 |
| *Pdzk1ip1* | 8.00 | 3.00 | 0.027 |
| *Pla1a* | 8.00 | 3.00 | 0.035 |
| *Csf3* | 7.87 | 2.98 | 0.029 |
| *Cblc* | 6.83 | 2.77 | 0.032 |
| *Spry4* | 6.44 | 2.69 | 0.029 |
| *Pf4* | 6.40 | 2.68 | 0.038 |
| *Jag2* | 6.17 | 2.62 | 0.038 |
| **Down-regulated** |  |  |  |
| *Ccl5* | -830.55 | -9.70 | 0.005 |
| *Serpinh1* | -261.33 | -8.03 | 0.008 |
| *Brca2* | -242.31 | -7.92 | 0.006 |
| *Dusp2* | -238.50 | -7.90 | 0.000 |
| *Tlr7* | -219.59 | -7.78 | 0.013 |
| *Arnt2* | -217.11 | -7.76 | 0.016 |
| *Pias4* | -180.14 | -7.49 | 0.005 |
| *Map3k12* | -145.56 | -7.19 | 0.022 |
| *P4ha1* | -100.22 | -6.65 | 0.035 |
| *Msh2* | -85.50 | -6.42 | 0.000 |
| *Bax* | -74.50 | -6.22 | 0.012 |
| *Tnfrsf4* | -74.50 | -6.22 | 0.000 |
| *Parp12* | -60.71 | -5.92 | 0.009 |
| *Hes1* | -60.24 | -5.91 | 0.019 |
| *Angptl4* | -59.50 | -5.89 | 0.001 |
| *Ctss* | -49.33 | -5.62 | 0.030 |
| *Bad* | -46.30 | -5.53 | 0.010 |
| *Inhba* | -35.26 | -5.14 | 0.001 |
| *Il34* | -33.93 | -5.08 | 0.010 |
| *Prr5* | -32.50 | -5.02 | 0.001 |

In the “CEI-20 vs. naïve” comparison, there were 56 significantly regulated TM genes, with 23 being up-regulated and 33 down-regulated (Figure 4). Table 3 shows 20 up-regulated and 20 down-regulated genes. A complete list of the 56 genes can be found in Supplementary Table S4. These 56 genes were deemed “cannulation-related”. Pathway analysis identified ‘cell activation’, ‘leukocyte activation’, ‘immune system development’, and ‘inflammatory response’ genes as top ‘GO biological process’ pathways, which is consistent with our hypothesis that this comparison would yield genes regulated in response to cannulation or anesthesia.

**Figure 4.**
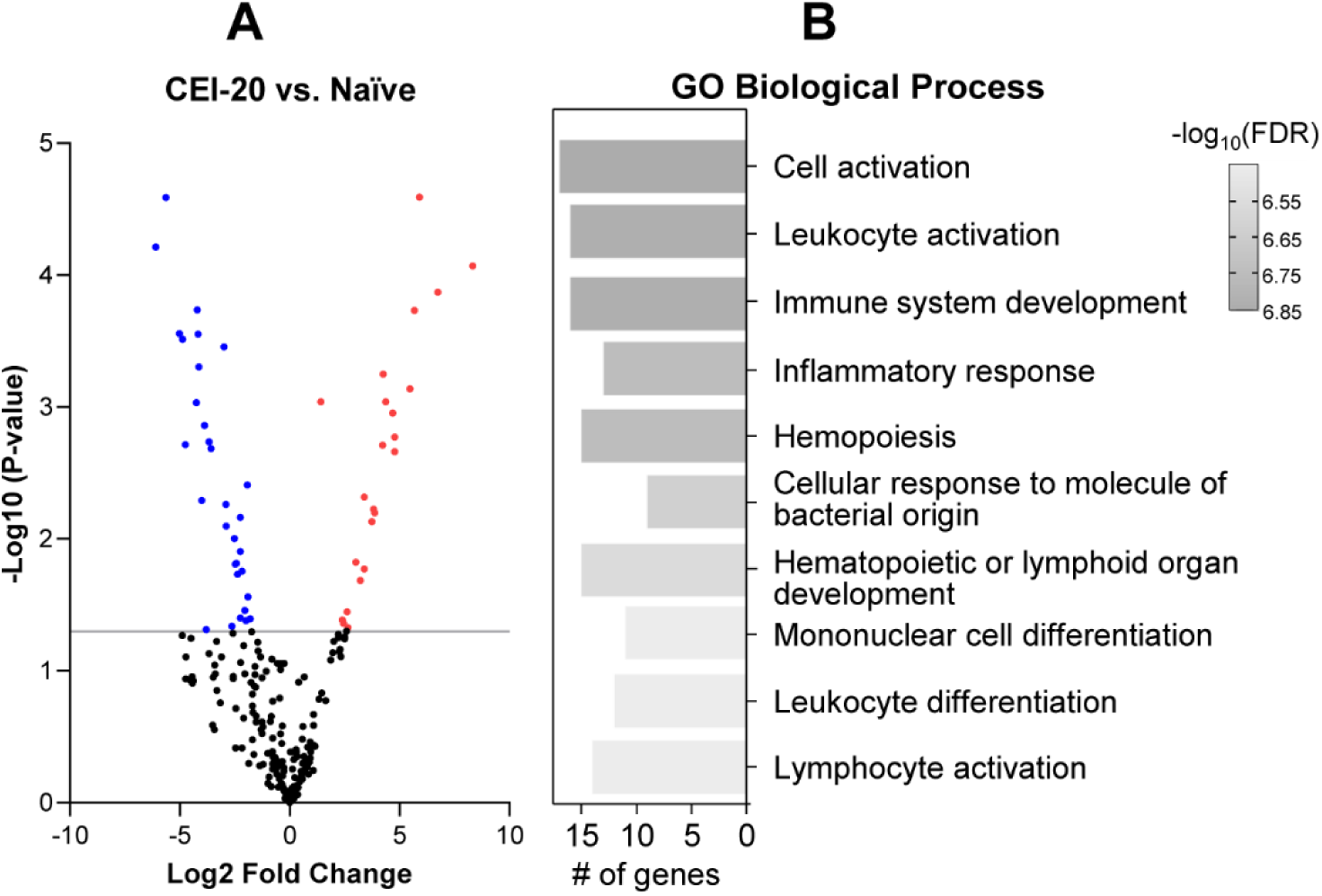
Nanostring analysis of CEI-20 versus naïve groups. (A) Volcano plot showing all genes on the cartridge. Significantly up-regulated genes are shown in red and down-regulated genes are shown in blue. (B) ShinyGO 0.77 analyses shows the top 10 most significantly affected ‘GO biological process’ pathways.

**Table 3:** Twenty up-regulated and 20 down-regulated TM genes significantly differentially expressed in “CEI-20 vs. naïve” (Fold change >1.5 or <-1.5; p-value <0.05).

| Name | Fold Change | Log2 FC | p-Value |
| --- | --- | --- | --- |
| **Up-regulated** |  |  |  |
| *Il6* | 321.62 | 8.33 | 8.52E-05 |
| *Birc3* | 106.25 | 6.73 | 1.35E-04 |
| *Tnfaip6* | 60.02 | 5.91 | 2.57E-05 |
| *Ccl2* | 50.91 | 5.67 | 1.85E-04 |
| *Fosl1* | 44.25 | 5.47 | 0.001 |
| *Clec4e* | 27.50 | 4.78 | 0.002 |
| *Mmp13* | 27.50 | 4.78 | 0.002 |
| *Csf3* | 25.85 | 4.69 | 0.001 |
| *Myc* | 20.58 | 4.36 | 0.001 |
| *Cebpb* | 19.11 | 4.26 | 0.001 |
| *Nfil3* | 18.63 | 4.22 | 0.002 |
| *Tnfaip3* | 14.63 | 3.87 | 0.006 |
| *Sell* | 14.13 | 3.82 | 0.006 |
| *Ripk3* | 13.30 | 3.73 | 0.007 |
| *Stc1* | 10.52 | 3.40 | 0.005 |
| *Ccl7* | 10.50 | 3.39 | 0.017 |
| *Pla1a* | 9.25 | 3.21 | 0.021 |
| *Tap1* | 8.07 | 3.01 | 0.015 |
| *Nlrp3* | 6.25 | 2.64 | 0.047 |
| *Pf4* | 6.10 | 2.61 | 0.036 |
| **Down-regulated** |  |  |  |
| *Msh2* | -68.40 | -6.10 | 6.12E-05 |
| *Tnfrsf4* | -49.67 | -5.63 | 2.58E-05 |
| *Prr5* | -32.50 | -5.02 | 2.77E-04 |
| *Cdkn1c* | -29.29 | -4.87 | 3.07E-04 |
| *Crabp2* | -26.85 | -4.75 | 0.002 |
| *Fgf18* | -19.00 | -4.25 | 0.001 |
| *Flt1* | -18.57 | -4.21 | 0.000 |
| *Hdac11* | -18.00 | -4.17 | 0.000 |
| *Mx1* | -17.50 | -4.13 | 0.000 |
| *Dusp2* | -16.03 | -4.00 | 0.005 |
| *Ube2t* | -14.75 | -3.88 | 0.001 |
| *Bad* | -13.88 | -3.80 | 0.049 |
| *Ccno* | -12.75 | -3.67 | 0.002 |
| *Eif4ebp1* | -12.75 | -3.67 | 0.002 |
| *Trem2* | -12.00 | -3.58 | 0.002 |
| *Sfxn1* | -8.00 | -3.00 | 0.000 |
| *Myd88* | -7.50 | -2.91 | 0.005 |
| *Sox11* | -7.40 | -2.89 | 0.008 |
| *Sbno2* | -6.20 | -2.63 | 0.046 |
| *Bnip3l* | -5.75 | -2.52 | 0.010 |

To identify IOP-responsive TM genes, we compared the 95 unique genes identified in the two hypertensive comparisons (“CEI-50 vs. CEI-20” and “CEI-50 vs. naïve”) to the list of 56 “cannulation-related” genes. This yielded a total of 151 genes, with 45 of these being only significant in the hypertensive comparisons. We therefore considered there to be 45 IOP-responsive genes. (Table 4; Figure 5). We analyzed the ‘GO cellular component’ pathways with these 45 IOP-responsive genes. Of the top 10 pathways identified, genes related to extracellular matrix (‘extracellular space’ and ‘extracellular region’) and plasma membrane (‘membrane raft’ and ‘membrane microdomain’) were significantly regulated (Figure 5A). Genes identified as being ECM-related were: *Angptl4, Ccl5, Ctss, Cxcl3, Ifi35, Il1a, Inhba, Mfge8, Prom1, Serpinh1,* and *Wnt7b*. Genes associated with the plasma membrane were: *Aldoa, Aplnr, Ccr9, Cd28, Ctnnb1, Fcgr2b, Ifitm2, Jag2, Jak2, Map3k12, Oas3, Tlr7,* and *Zap70*.

**Figure 5.**
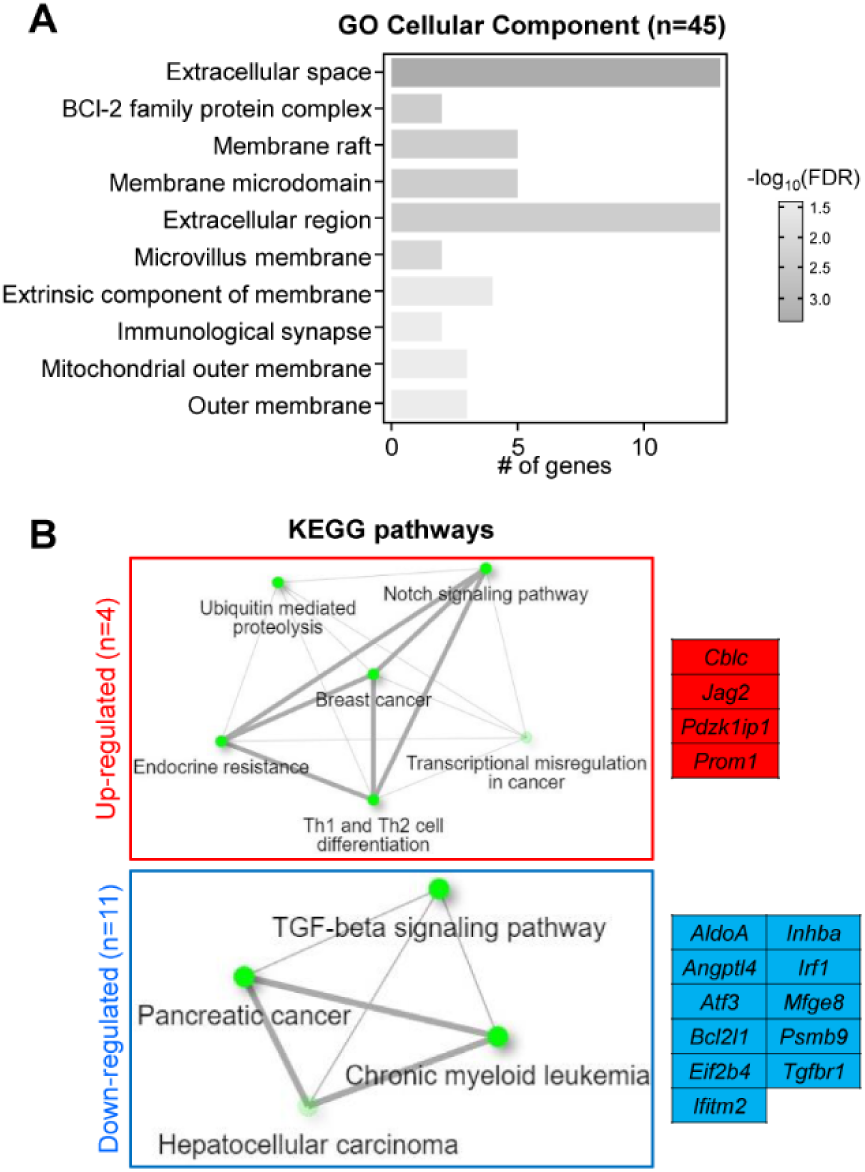
Pathway analysis of 45 IOP-related genes. (A) ShinyGO 0.77 analyses shows the top 10 most significantly affected ‘GO cellular component’ pathways. (B) KEGG pathway analysis of the 4 up-regulated and 11 down-regulated genes that were common to the “CEI-50 vs. CEI-20”, and “CEI-50 vs. naïve” groups

**Table 4.**
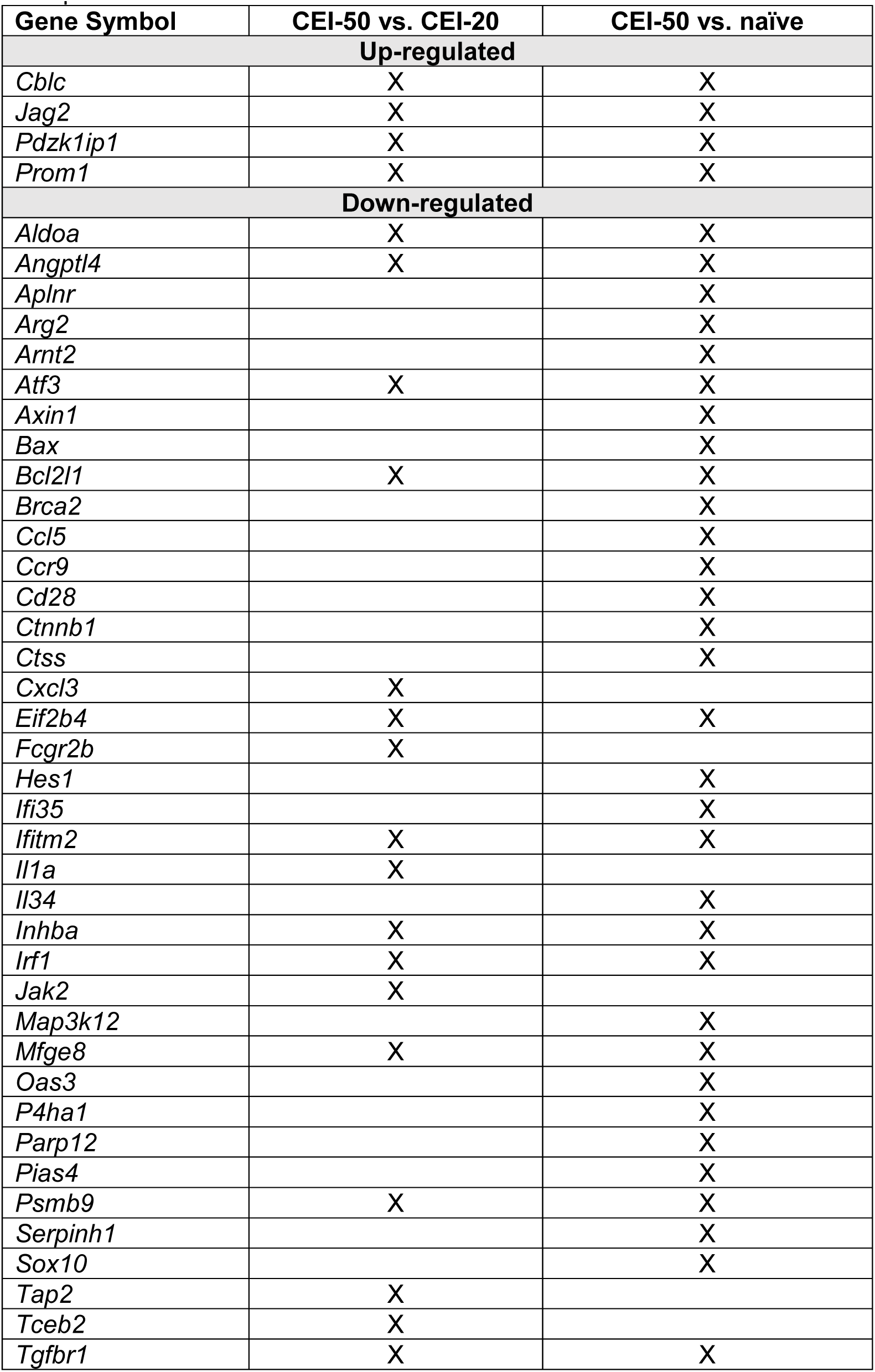
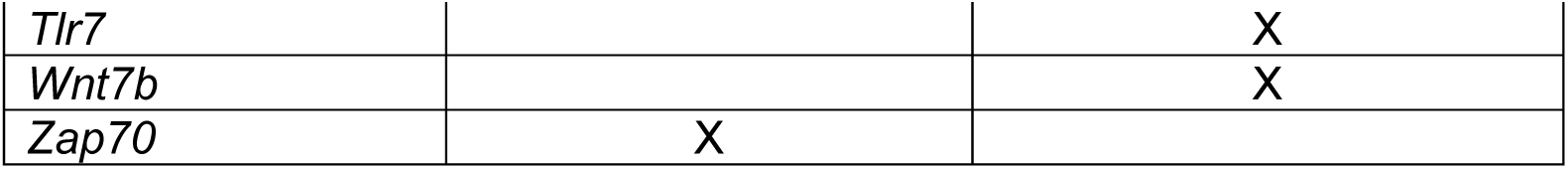
All IOP-related TM genes (n=45). TM genes significantly differentially expressed in “CEI-50 vs. CEI-20” and “CEI-50 vs. naïve” comparisons, but without those genes common to “CEI-20 vs. naïve” comparison.

Of these 45 IOP-related genes, 15 genes were regulated in the same direction in both hypertensive comparisons. Four genes were up-regulated (*Cblc, Jag2, Pdxk1IP1*, *Prom1),* and 11 were down-regulated (*Aldoa, Angpl4, Atf3, Bcl2l1, Eif2b4, Ifitm2, Inhba, Irf1, Mfge8, Psmb9, Tgfbr1*) (Figure 5B). KEGG pathway analysis showed that ‘Notch signaling’ and ‘ubiquitin-mediated proteolysis’ were affected by up-regulated genes, while the TGFβ pathway was significantly affected by down-regulated genes (Figure 5B). Normalized counts for all 750 genes for each sample in the TM groups are found in Supplemental Table S5.

### ONH gene analysis by Nanostring

We also analyzed ONH RNA from tissue dissected from the same group of hypertensive rats described above. In the “CEI-50 vs. naïve” comparison, we identified 22 significantly regulated genes (Table 5). Several of these genes (*Angptl4, Cebpb, Edn1, Myd88, Nfil3, Slc2a1,* and *Srebf1*) were regulated in the same direction between this Nanostring study and our published ONH RNA-seq study (Table 6). ^27^ Pathway analysis identified ‘immune response’, ‘response to cytokine’ and ‘response to interleukin-1’ as examples of significantly affected biological process pathways immediately after CEI (Figure 6).

**Figure 6.**
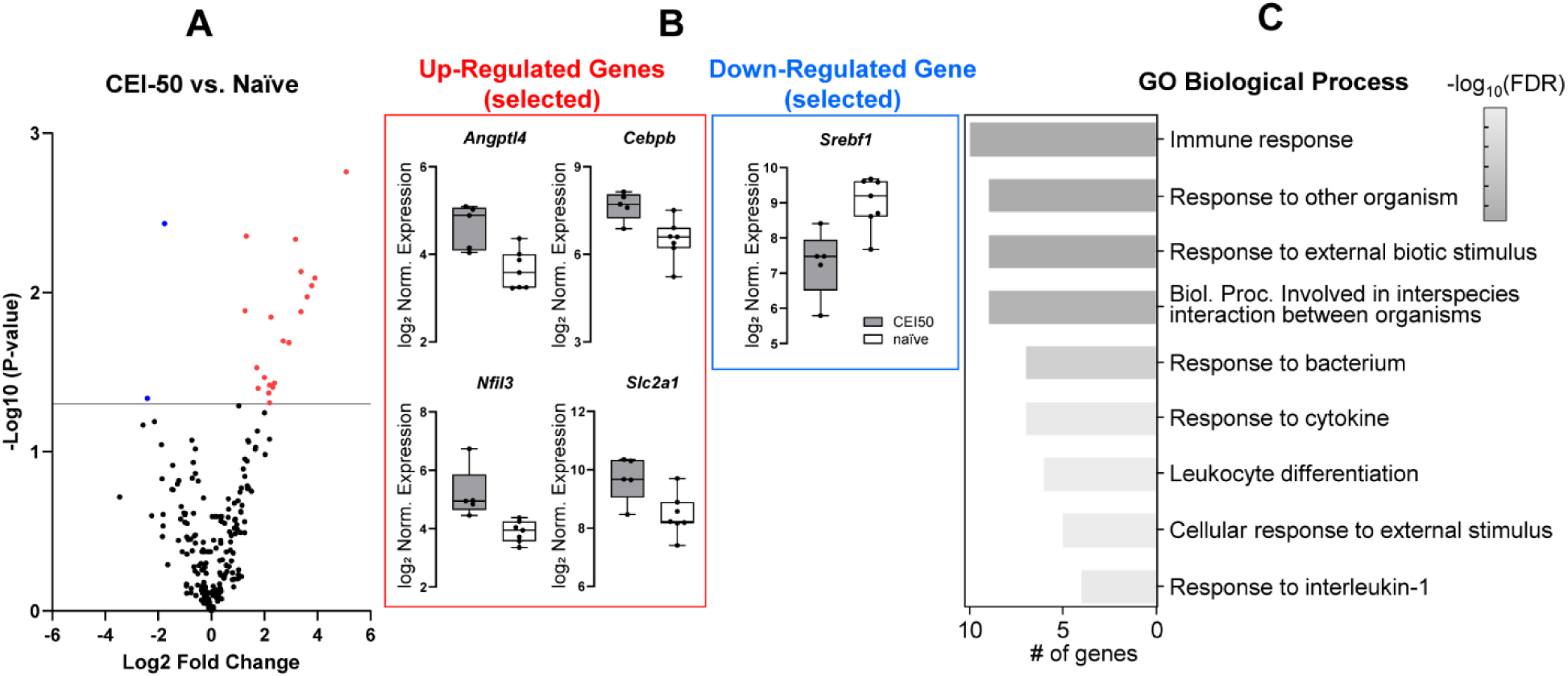
Nanostring analysis of optic nerve head CEI-50 versus naïve groups. (A) Volcano plot showing all genes on the cartridge. Significantly up-regulated genes are shown in red and down-regulated genes are shown in blue. (B) Normalized expression levels of selected up- and down-regulated genes in CEI-50 (n=5; grey) and naïve (n=7; white) are shown. (C) ShinyGO 0.77 analyses shows the top most significantly affected ‘GO biological process’ pathways.

**Table 5:** All ONH genes differentially expressed in “CEI-50 vs. naïve” (Fold change >1.5 or <-1.5; p-value <0.05).

| Gene Symbol | Fold Change | Log2 FC | p-Value |
| --- | --- | --- | --- |
| **Up-regulated** |  |  |  |
| *Myd88* | 34.02 | 5.09 | 0.002 |
| *Irf4* | 15.00 | 3.91 | 0.008 |
| *Il12rb2* | 13.80 | 3.79 | 0.009 |
| *Parp4* | 12.20 | 3.61 | 0.011 |
| *Nfil3* | 10.42 | 3.38 | 0.007 |
| *Tnfsf8* | 10.40 | 3.38 | 0.013 |
| *Edn1* | 9.00 | 3.17 | 0.005 |
| *Fcgrt* | 7.60 | 2.93 | 0.021 |
| *H2-DMa* | 6.53 | 2.71 | 0.020 |
| *Fasl* | 5.20 | 2.38 | 0.037 |
| *Itga6* | 5.00 | 2.32 | 0.039 |
| *Cd163* | 4.76 | 2.25 | 0.014 |
| *Got1* | 4.59 | 2.20 | 0.038 |
| *Snca* | 4.59 | 2.20 | 0.049 |
| *Angptl4* | 4.48 | 2.16 | 0.043 |
| *Tnfrsf11a* | 4.01 | 2.00 | 0.034 |
| *Clec4e* | 3.40 | 1.77 | 0.040 |
| *C7* | 3.27 | 1.71 | 0.030 |
| *Cebpb* | 2.50 | 1.32 | 0.004 |
| *Slc2a1* | 2.42 | 1.27 | 0.013 |
| **Down-regulated** |  |  |  |
| *Serpinh1* | -5.32 | -2.41 | 0.046 |
| *Srebf1* | -3.40 | -1.77 | 0.004 |

**Table 6.**
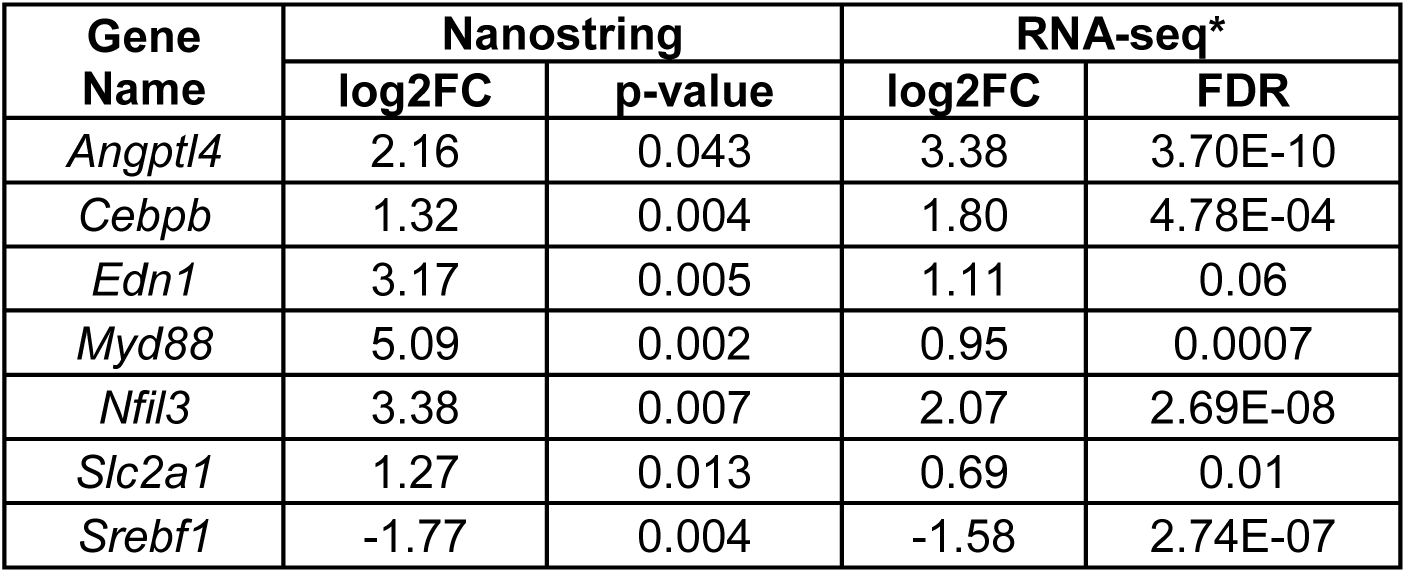
ONH genes identified by both Nanostring and RNAseq following an 8-hour pressure exposure. *, see reference [27].

## Discussion

Elevated IOP is a known risk factor for glaucoma. IOP becomes elevated due to obstructed aqueous humor outflow pathway in the anterior chamber leading to optic nerve degeneration in the posterior pole. Yet, most animal models of glaucoma cause damage to the TM, causing IOP to become elevated, but render the TM unusable for functional analyses. To overcome this, we used the CEI rat model to study *in vivo* TM gene responses to elevated IOP because the cannulation tube does not damage the TM. This is the first study to identify endogenous TM IOP-responsive genes *in vivo*. Furthermore, we report TM and ONH gene responses from the same cohort of animals.

In response to a pressure challenge of 50 mmHg, approximately 2.5x mean rat daytime IOP, we detected 45 IOP-responsive genes in the TM. Gene ontology pathway analysis highlighted 13 genes in ECM-related pathways to be significantly regulated. This is consistent with *ex vivo* perfusion studies, and cell mechanical stretch experiments, which have led to the hypothesis that ECM remodeling adjusts the TM outflow resistance as a pressure-induced homeostatic response.^1,2,6,8^

We compared the 45 IOP-related genes identified in this *in vivo* rat IOP study to those genes identified in pressure-challenged human perfusion culture studies, ^2, 12, 13^ or with TM cells subject to mechanical stretch (Table 7). ^6, 7^ While there are differences in species studied (rat, human, pig), experimental design (pressure level and duration), and assay method (microarray, PCR arrays, Nanostring), *ANGPTL4* was detected among all studies. ANGPTL4 is a secreted, matricellular protein, which can bind cell surface integrins and fibronectin. Interestingly, interaction of ANGPTL4 with fibronectin delays its proteolytic degradation by MMPs. ^38^ In the rat TM, *Angptl4* was down-regulated, implying that proteolytic sites in fibronectin may become exposed and more readily available for IOP-induced MMP degradation. In addition, studies with retinal vascular endothelial cells have implicated *ANGPTL4* in activating the Rho/ROCK signaling pathway, which controls actin cytoskeleton contractility. ^39^ This suggests that reduced *Angptl4* gene expression would suppress Rho/ROCK activity, leading to relaxation of TM cells, increased aqueous outflow, and IOP reduction.

**Table 7.** Comparison of 45 rat TM IOP-related genes compared to genes identified to be significantly altered in pressure-challenged human perfusion cultures, or TM cell mechanical stretch.

| **Study Lead Author** | **Method** | **Species** | **IOP and duration** | **Assay method** | **Total # genes in the study** | **Genes in common with this *in vivo* study** | **Ref.** |
| --- | --- | --- | --- | --- | --- | --- | --- |
| **Gonzalez** | Perfusion | Human | 50 mmHg, 6 hours | Microarray | 8 | - | #12 |
| **Vittitow** | Perfusion | Human | 50 mmHg, 48 hours | Microarray | 40 | - | #2 |
| **Vittal** | Cell mechanical stretch | Pig | Static stretch 12, 24, 48 hours | Microarray | 143 | *CCL7*<br>*IRF1* | #6 |
| **Luna** | Cell mechanical stretch | Human | 15% stretch, 1 cycle/sec, 6 hours | Microarray | 152 | *ANGPTL4*,<br>*CCL7*<br>*IRF1* | #7 |
| **Vranka** | Perfusion | Human | 16 mmHg, 48 hours | PCR array | 34 | *CTTNB1* | #13 |

Several secreted molecules were significantly regulated in the TM, including up-regulation of *Jag2*, and down-regulation of *Mfge8* and *Wnt7b*. Jagged-2 (*Jag2*) is a ligand for the Notch family of receptors. ^40^ Notch signaling is important for cellular communication between adjacent cells and may play important functional roles in mechanotransduction. ^41^ Upon binding of the JAG2 ligand, Notch receptor is proteolytically cleaved both extracellularly and intracellularly, with the intracellular fragment translocating to the nucleus to activate or inhibit transcription. Thus, Notch may act as a ‘mechanostat’ that maintains the stasis of tissues subject to mechanoforces. ^41^ Up-regulation of *Jag2* is consistent with this paradigm. Milk fat globule EGF-factor-8 (*Mfge8*), also known as lactadherin, is a secreted glycoprotein that binds to integrin β3 and β5 chains, and is dexamethasone responsive in TM cells. ^42^ It functions as an ‘eat me’ signal, being up-regulated on apoptotic or injured cells, thereby labeling them for phagocytosis. *Mfge8* is also an abundant component of TM-derived exosomes. ^43^ *Wnt7b* is a short-range signaling protein involved in canonical Wnt signaling via Frizzled receptors. The canonical Wnt signaling pathway regulates IOP and ECM synthesis in the TM. ^20, 44^ Down-regulation of Wnt ligands, such as *Wnt7b* in this study, results in intracellular degradation of β-catenin, which is therefore unable to translocate to the nucleus and transcription of Wnt target genes are therefore suppressed. ^45^ Interestingly, the β-catenin (*Ctnn1*) gene is also down-regulated in the IOP-challenged rat TM.

Several transmembrane receptors were identified as significantly altered by pressure challenge. including *Fcgr2b*. Clustering of the *Fcgr2b* receptor induces phagocytosis and synthesis of inflammatory cytokines. ^46^ Down-regulation of *Fcgr2b* in pressure-challenged TM suggests that these cellular functions are suppressed during IOP homeostasis. Surprisingly, only one of the 12 integrins on the panel was significantly different in the hypertensive group comparisons. This was *Itgae* (-4.75 fold change; -2.25 Log_2_FC in the “CEI-50 vs. naïve” comparison). *Itga6* was significantly different in the CEI-20 versus naïve comparison. Integrins are transmembrane proteins that detect changes to the biomechanical environment, such as stretch/distortion resulting from elevated IOP. ^47^ Despite the lack of significant changes at the gene level, mechanical stretch induces integrin proteins to undergo conformational changes that lead to structural activation, which propagates biomechanical signals intracellularly. Thus, even if integrin gene expression is unchanged, it is likely that integrin-mediated signaling occurs in response to pressure challenge.

Analysis of genes common to both hypertensive comparisons revealed that the TGFβ pathway was significantly regulated. The TGFβ pathway has been closely studied in relation to glaucoma since the first report of increased levels of TGFβ2 in the aqueous humor of POAG patients. ^48^ Increased TGFβ2 induces ECM protein synthesis, thereby contributing to the fibrotic-like matrix characteristic of the glaucomatous TM. ^49^ Of the TGFβ pathway genes on the panel in this study, Pdzk1-interacting protein-1 (*Pdzk1ip1*) was up-regulated, while TGFβ-receptor-1 (*Tgfbr1*) and inhibin-beta A (*Inhba*) were significantly down-regulated in both TM hypertensive comparisons. *Pdzk1ip1* interacts with intracellular target of TGFβ signaling, SMAD4, thereby preventing translocation of SMADs from the cytosol to the nucleus. Thus, *Pdzk1ip1* is antagonist of TGFβ signaling. *Inhba* encodes activin beta, a ligand for the TGFβR1. Once activin engages its receptor, SMAD2/3 proteins become phosphorylated, initiating SMAD4 translocation to the nucleus, which then modulates expression of many target genes. ^50^ Reduced synthesis of a TGFβ ligand and receptor, with concomitant up-regulation of a TGFβ antagonist, suggests that elevated IOP prevents activation of this signaling pathway, thereby preventing ECM over-production. Thus, our results suggest that an *in vivo* 8-hour pressure challenge induces protective pathways that induce homeostasis, but prevent TGFβ-induced TM fibrosis.

There were 56 TM genes identified in the “CEI-20 vs. naïve” comparison that were deemed ‘cannulation-related’. A previous study showed that transcorneal needle puncture significantly lowered IOP at 24 hours following puncture (Schuman et al., ARVO 2023 annual meeting abstract, 2048). We would not be able to detect similar IOP decreases caused by needle puncture placement since we precisely control the pressure challenge. Transcorneal needle puncture also significantly increased macrophage density of distal outflow vessels (Schuman et al., ARVO 2023 annual meeting abstract, 2048). Our cannulation procedure is likely to induce similar inflammation-related events in the anterior chamber, and perhaps into the distal outflow pathway. When the 56 genes were analyzed, IL-17, TNFα, and NOD-like KEGG pathways, and the GO biological process ‘inflammatory response’ were significantly affected. These are consistent with inflammation. However, several of these TM ‘cannulation-related’ genes can directly affect IOP. For instance, *Rock1* is a major effector of the actin cytoskeleton, and a target for Rho kinase inhibitors, the newest class of glaucoma drugs. ^51^ Down-regulation of Rock1 in this comparison is consistent with TM relaxation and increased aqueous outflow. Another gene affected in the “CEI-20 vs. naïve” comparison was *Vegfa,* which was up-regulated. VEGFa protein is secreted by TM cells in response to mechanical stretch and application of exogenous VEGFa in enucleated mouse eyes increased outflow facility. ^52^ Thus, while we discounted these genes from our IOP-related genes, our somewhat simplistic approach of subtracting one set of genes from another gene list may have discarded some genes that regulate IOP *in vivo*.

In a previous study, we exposed animals to a CEI of 20 or 60 mmHg for 8-hours and analyzed the ONH transcriptome by RNA-sequencing. ^27^ ONH were collected 0hr and 1 to 10 days following CEI and compared to ONH from naïve animals. Following CEI of 20 mmHg, there were only 18 significantly regulated ONH genes at 0hr and none at the other time points. These 18 genes did not cluster into any significant gene category and were not significant following a CEI of 60 mmHg. By comparison, we identified 1,354 significantly regulated genes immediately following (0hr) CEI of 60 mmHg, the same time-point as used in the current study. These findings support that ONH gene responses following a CEI of 60 mmHg are responses to elevated IOP and not a by-product of anesthesia or anterior chamber cannulation performed during CEI. The top gene ontology category immediately (0hr) following CEI of 60 mmHg was defense response. These 0hr time point genes likely play an early role in glaucomatous neurodegeneration that may be important regulators of downstream cellular events. For example, we previously identified that cellular proliferation of astrocytes is an early cellular event following chronic and transient elevations of IOP. ^37, 53, 54^ From our RNA-seq study, we were able to identify several key genes that reflect signaling or regulation of ONH cellular proliferation at 0hr. Importantly, the highest number of significantly regulated ONH genes occurred immediately following CEI (0hr), relative to the other time points analyzed (1-10 days). Therefore, we decided to re-analyze this early time point by Nanostring technology but following a CEI of 50 mmHg, 10 mmHg lower that our previous RNA-Seq study.

In spite of the lower pressure exposure, we were able to identify 22 significantly regulated ONH genes by Nanostring. More than half of these 22 genes are involved with an immune response (*C7, Cebpb, Clec4e, Edn1, Fasl, H2-DMa, Irf4, Myd88, Nfil3, Snca, Srebf1*, *Tnfrsf11a,* and *Tnfsf8*). Despite the different levels of IOP, there were seven genes in common between the current Nanostring data and our published RNA-seq findings (Table 6). ^27^ Three genes (*Angptl4, Cebpb, Slc2a1*) have previously been reported to be upregulated and secreted by cortical astrocytes in Alzheimer’s and Parkinson’s disease. ^55–59^ In addition, endothelin-1 (*Edn1*) is significantly elevated in glaucomatous optic nerves, where it induces astrocyte proliferation, ^60^ and negatively affects blood flow and retinal ganglion cell viability. ^61, 62^ It is important to highlight that out of the 750 genes in the Nanostring panel, only 109 of these were significant in our previous RNA-seq study. In spite of the lower number of significantly altered ONH genes identified by Nanostring, the most prominent gene category remained related to an immune/defense response, as previously identified by our RNA-seq study. The lower number of significantly regulated genes identified by Nanostring could have been influenced by technological differences, the lower IOP level used in the current study, and/or selecting a panel that was more specific for TM cellular events. In this study, RNA was extracted from an ONH region that extended from Bruch’s membrane to 400µm posterior, a region that is enriched with Sox2+ nuclei, an astrocytic marker, suggesting that astrocytes are the predominant cell type in this ONH region. ^53^ Therefore, it is likely that these genes are also expressed/secreted by ONH astrocytes as well. These findings are consistent with astrocytes playing a key role in the early response to elevated pressure.

A major advantage of our approach is the ability to assay gene responses in both TM and ONH simultaneously. As expected, the heatmap in Figure 1, generated from naïve datasets, shows that the two tissues express unique gene sets. This is not surprising because during development, the retina and optic nerve arise from neuroectoderm, while TM develops from mesenchymal neural crest cells. ^63, 64^ Two genes, *SerpinH1* and *Angptl4*, were differentially regulated by elevated IOP in both TM and ONH. *SerpinH1* was down-regulated in both tissues. *SerpinH1* (heat shock protein 47) is an important chaperone for fibrillar collagen biosynthesis. It has a role in fibrosis in several tissues, and down-regulation would be consistent with decreased fibrosis. ^65^ On the other hand, *Angptl4* was up-regulated in ONH, but down-regulated in TM suggesting it may induce different downstream effectors in each tissue.

This is the first study to compare directly the responses of the TM and ONH within the same animal to a given level of IOP. It is extremely interesting that, immediately following an 8-hour pressure exposure, the TM demonstrates a range of responses encompassing several different pathways, many of which agree with published *in vitro* work. The robust nature of this response so soon after a pressure challenge is entirely consistent with the notion that maintaining homeostasis a fundamental function of the TM. This is important for overall eye stability as well as possibly protecting the TM itself. While relatively fewer regulated genes were identified in the ONH, they represent pathways that largely mirror “defense response” identified at 0hr in our earlier RNA-seq study. As with the TM, it is reasonable to hypothesize that these early gene responses may also be protective. We believe the CEI model, which reveals a sequence of cellular events following a single pressure challenge, can be used to test this hypothesis and allow us to better understand the relationship of these early ONH changes to axonal injury.

There are several limitations of this study. First, the panel only contained 750 genes and did not include several known IOP-responsive genes important for remodeling TM ECM, such as MMP2, MMP3, and MMP14. ^8, 66^ In addition, many genes identified in the rat ONH RNA-seq study were not on the Nanostring panel. Thus, our study is likely missing some key genes that respond to IOP *in vivo*. Also, the predesigned Nanostring panel is designed for mouse genes. While there is generally good homology between rat and mouse DNA sequences, we found that 17% of these mouse genes were less than 80% identical to the target rat gene, possibly impacting the detection of rat RNA on the panel. Nevertheless, we detected significant changes to 45 IOP-related genes in the TM, and 22 differentially expressed genes in the ONH. TaqMan qPCR analysis using rat-specific primers showed similar results to the Nanostring for several TM and ONH genes (Supplementary Figure S1), increasing our confidence that the gene changes detected by Nanostring are not artifactual. An additional limitation is that we used p-values to identify significantly regulated genes, not an adjusted p-value. However, the agreement between our Nanostring data, qPCR, and previous RNA-seq study, provide additional evidence that our findings are true representations of TM and ONH responses to elevated IOP. Finally, because animals cannot tolerate longer anesthesia, 8 hours is the maximum exposure duration that we assayed.

In summary, this study is the first to report TM and ONH IOP-responsive genes in the same animals. Our results indicate that in the TM, genes related to ECM synthesis and actin cytoskeleton are predominantly regulated *in vivo*, confirming *in vitro* and *ex vivo* studies that have implicated these pathways in IOP homeostasis. This study additional confirmed the early upregulation in immune/defense response-related genes, as previously identified by our RNA-seq study. The CEI model provides an excellent animal model system to study IOP responsive genes in both tissues that are involved in glaucoma.

## Acknowledgements

This study was supported by NIH/NEI grants R21 EY033073, R01 EY019643 (KEK), R01 EY032590 (KEK), R01 EY010145-17S1 (DCL), R01 EY010145 (JCM), P30 EY010572 (OHSU), the Malcolm M. Marquis, MD Endowed Fund for Innovation, and by unrestricted departmental funding from Research to Prevent Blindness (New York, NY).

---

**Supplementary Figure S1.**
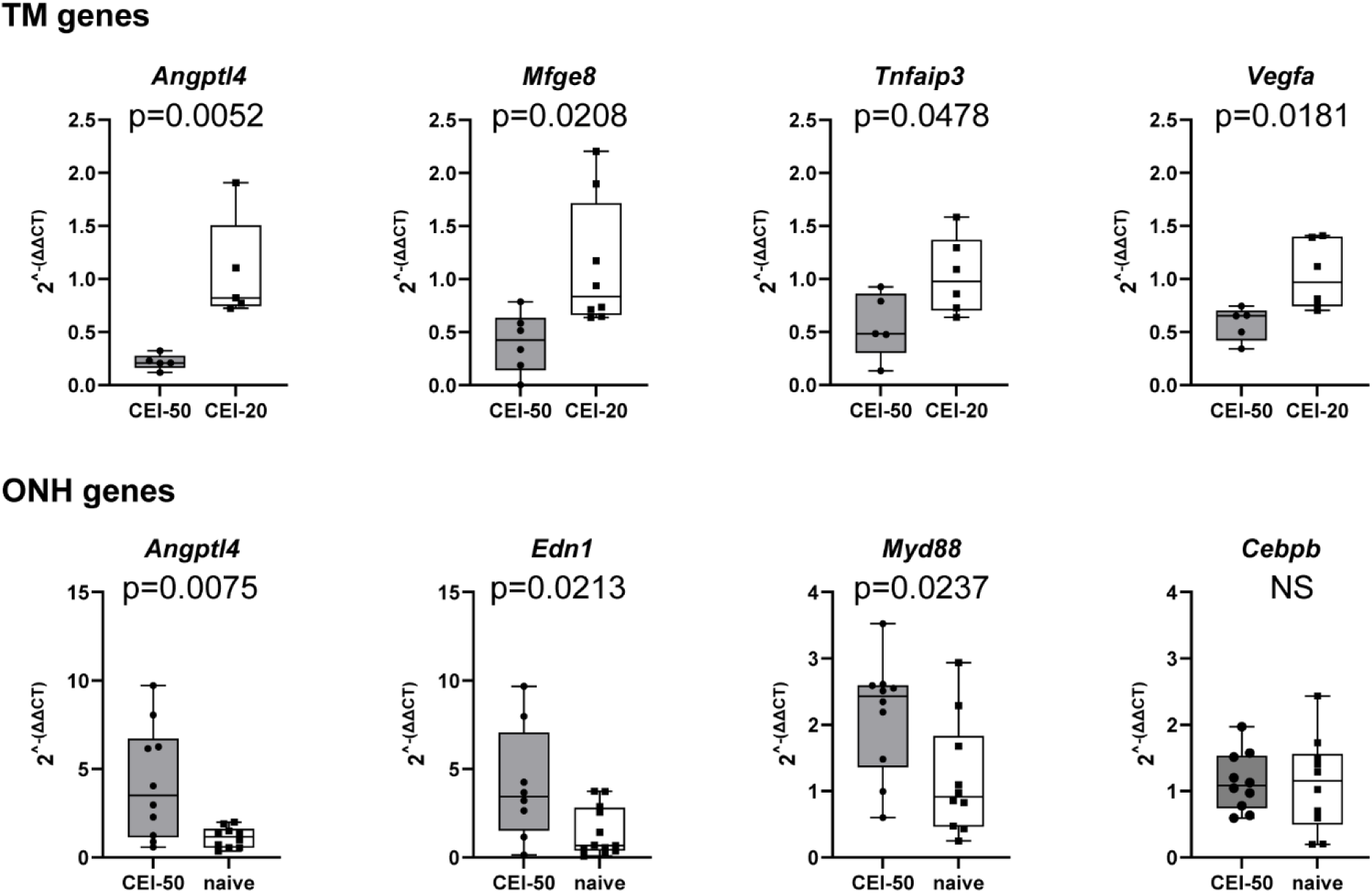
Taqman quantitative PCR analyses of selected TM and ONH genes. Data points represent biological and technical replicates. P-values were calculated by an unpaired t-test.

